# Disrupting the biodiversity – ecosystem function relationship: response of shredders and leaf breakdown to urbanization in Andean streams

**DOI:** 10.1101/2020.07.28.223867

**Authors:** Wilson Zúñiga-Sarango, Fernando P. Gaona, Valeria Reyes-Castillo, Carlos Iñiguez-Armijos

## Abstract

Urbanization is a major driver of stream ecosystems impairment and often associated with multiple stressors and species loss. A challenge is to understand how those stressors alter the relationship between biodiversity and ecosystem functioning (B-EF). In Andean streams of southern Ecuador, we assessed the response of shredder diversity and organic matter breakdown (OMB) to urbanization and identified the urban-associated stressors disrupting the B-EF relationship. A leaf-litter bag experiment during stable flow conditions in 2016 was carried out to quantify total OMB and shredder-mediated OMB, which was estimated to represent the B-EF relationship. We calculated the taxonomic and functional diversity of shredder invertebrates associated with leaf packs. Also, a suite of physicochemical and habitat stressors was weekly measured during the field experiment. Along the urbanization gradient, both taxonomic and functional diversity of shredders declined while OMB rates decayed. Shredders were absent and their contribution to OMB was null at the most urbanized sites. The B-EF relationship was interrupted through nutrient enrichment and physical habitat homogenization as a consequence of urbanization. These results demonstrate how species loss propagates to ecosystem functions in urbanized streams and how environmental stressors alter the B-EF relationship. Better land-use practices are crucial in Andean catchments to guarantee ecosystem services which are the result of a successful B-EF relationships.

## 1 Introduction

Aquatic ecosystems contain high biodiversity, but at the same time, they evidence a faster species loss than marine or terrestrial ecosystems (Collen et al., 2014; Sánchez-Bayo and Wyckhuys, 2019). Unfortunately, species loss also implies a reduction of ecosystem functions (Cardinale et al., 2006). Species diversity is crucial for the dynamics and functioning of ecosystems, for that reason the relationship between biodiversity and ecosystem functioning (B-EF) has been well established and is a central research issue (Loreau et al., 2001; Tilman et al., 2014). Also, the B-EF relationships can be altered by environmental gradients, complicating the prediction of the effects of species loss in ecosystems subjected to environmental stress (McKie et al., 2009). Therefore, it is imperative to understand the mechanisms that interrupt this relationship trough environmental disturbance gradients such as urbanization.

Urbanization is a complex process involving permanent changes in the landscape such as the transformation of land use from rural (i.e. forest, agriculture) to urban (Antrop, 2004). Also, urbanization is related to numerous stressors that interact synergistically on streams (Walsh et al., 2005), affecting the hydrology, chemistry and aquatic communities via multiple pathways (Paul and Meyer, 2001; Walsh et al., 2005). Urbanized streams often show a reduction in channel width and substrate size of streambed due to sediment inputs, as well as an increase in nutrient concentrations and other pollutants from sewage inputs and runoff (Allan, 2004). When these stream characteristics are altered by urbanization there will be a consequence on the macroinvertebrate communities (Roy et al., 2003; de Jesús-Crespo, Rebeca Ramírez, 2011; Hassett et al., 2018), resulting in species loss (Chadwick et al., 2006; Urban et al., 2006) that can lead to a disruption of the biodiversity–ecosystem functioning relationship (Meyer et al., 2005; Tilman et al., 2014). Therefore, the interruption of this interaction can turn into a serious issue at the ecosystem level when keystone species are removed or replaced from streams (Iñiguez-Armijos et al., 2016).

Macroinvertebrates can play important roles in stream ecosystem functioning (Wallace and Webster, 1996). For instance, organic matter breakdown (OMB) is mediated by shredder invertebrates, as well as by leaching, microbial conditioning, and physical abrasion (Gessner et al., 1999). Indeed, shredders importance on OMB has been well documented in different regions (Jonsson et al., 2001; Li and Dudgeon, 2009; Dangles et al., 2011). Shredders fragment coarse particulate organic matter (CPOM) into smaller particles to make organic carbon available for other aquatic consumers or ecosystem processes (Gessner et al., 1999; Ramírez and Gutiérrez-Fonseca, 2014). Most of leaf-shredding invertebrates in Neotropical streams belong to Trichoptera and Plecoptera, and a few taxa are members of Coleoptera, Ephemeroptera and Diptera (see Table 1 in Ramírez and Gutiérrez-Fonseca, 2014). Indeed, their presence is essential for the functioning of montane streams in the Neotropical region (Encalada et al., 2010; Dangles et al., 2011; Iñiguez-Armijos et al., 2016, 2018b), regardless of shredder diversity can be lower here compared to temperate streams (Boyero et al., 2011). Unfortunately, as an effect of urbanization, most of the sensitive aquatic insects (i.e. Trichoptera, Plecoptera, Ephemeroptera) are reaching alarming rates of species loss worldwide (Sánchez-Bayo and Wyckhuys, 2019) removing taxa with key functional feeding habits (e.g. Trichoptera shredders), “breaking down” the linkages of these taxa with crucial ecosystem functions such as OMB.

**Table 1.** Set of the environmental variables, shredder diversity and breakdown rates measured in Andean streams (n = 12) along an urbanization gradient in southern Ecuador. The symbol ‡ indicates the variable significantly loaded to an axis of the sparse Principal Components Analysis.

|  | Units | Mean | Min | Max | PC1 | PC2 |
| --- | --- | --- | --- | --- | --- | --- |
| **Water chemistry** |  |  |  |  |  |  |
| Temperature | °C | 15.8 | 13.9 | 17.7 |  |  |
| Conductivity | µS·cm<sup>-1</sup> | 118.7 | 10.0 | 374.0 |  |  |
| pH |  | 7.9 | 7.1 | 8.6 | ‡ |  |
| DO | mg·L<sup>-1</sup> | 6.8 | 2.2 | 8.3 | ‡ |  |
| DIN | mg·L<sup>-1</sup> | 4.1 | 0.3 | 19.2 | ‡ |  |
| DRP | mg·L<sup>-1</sup> | 0.5 | 0.1 | 2.6 | ‡ |  |
| Turbidity | NTU | 50.9 | 0.6 | 320.0 | ‡ |  |
| **Hydraulic** |  |  |  |  |  |  |
| Channel width | m | 2.8 | 1.0 | 6.4 | ‡ |  |
| Channel depth | cm | 16.2 | 5.8 | 43.5 | ‡ |  |
| Current velocity | cm·s<sup>-1</sup> | 0.5 | 0.2 | 1.2 | ‡ |  |
| **Substrate size** |  |  |  |  |  |  |
| Silt | % | 15.7 | 0.0 | 75.0 |  |  |
| Sand | % | 10.0 | 0.0 | 32.5 |  |  |
| Gravel | % | 21.9 | 0.0 | 55.0 |  |  |
| Pebble | % | 23.8 | 5.0 | 67.5 |  |  |
| Cobble | % | 16.9 | 0.0 | 45.0 |  |  |
| Boulder | % | 11.5 | 0.0 | 40.0 |  |  |
| Bedrock | % | 0.0 | 0.0 | 0.0 |  |  |
| Substrate index |  | 29.7 | 9.7 | 87.7 | ‡ |  |
| **Land cover** |  |  |  |  |  |  |
| Canopy cover | % | 53.3 | 16.6 | 82.6 |  | ‡ |
| Forest | % | 49.0 | 11.8 | 99.2 |  |  |
| Pasture | % | 42.0 | 0.8 | 75.3 |  |  |
| Urban | % | 8.7 | 0.0 | 43.3 | ‡ |  |
| **Shredder invertebrates** |  |  |  |  |  |  |
| Richness | taxa/bag<sup>-1</sup> | 1.6 | 0.0 | 6.0 |  |  |
| Abundance | ind./bag<sup>-1</sup> | 5.9 | 0.0 | 52.0 |  |  |
| **Breakdown rates** |  |  |  |  |  |  |
| *k*<sub>coarse</sub> | dd<sup>-1</sup> | 0.0019 | 0.0001 | 0.0056 |  |  |
| *k*<sub>fine</sub> | dd<sup>-1</sup> | 0.0014 | 0.0007 | 0.0043 |  |  |
| *k*<sub>shredders</sub> | dd<sup>-1</sup> | 0.0004 | -0.0011 | 0.0041 |  |  |

Ecosystem functioning refers to ecological processes determined by the activity of biodiversity (Tilman et al., 2014). Nonetheless, this close relationship can be altered by several environmental stressors. In accordance with the Millennium Ecosystem Assessment (MEA, 2005), urbanization has a large effect on the integrity of freshwater ecosystems and its associated environmental stressors (e.g. nutrient enrichment) are affecting seriously stream ecosystem functions such as OMB (Meyer et al., 2005). Naturally, nutrients have a great role in OMB in the tropics (Tiegs et al., 2019), and a nutrient enrichment will disturb this ecosystem function as well as other stream parameters. Several studies have reported that urbanization has negatively affected both decomposer communities and OMB by nutrient enrichment and hydrologic alteration (e.g. Chadwick et al., 2006; Martins et al., 2015; Iñiguez-Armijos et al., 2016). However, because biodiversity and ecosystem functioning respond to stressors in different pathways (Sandin and Solimini, 2009), the joint application of structural and functional metrics to quantify the response of stream ecosystems to a stressor such as urbanization is adequate (Clapcott et al., 2012). Taxonomic metrics can indicate how the organization of aquatic communities responds to environmental stress, while functional biological traits can provide information on multiple-stressor effects and insights into potential mechanisms (Lange et al., 2014). Besides, if ecosystem processes such as OMB rates are quantified, a more comprehensive response to environmental stress gradients can be obtained (Riipinen et al., 2009; Voß et al., 2015), which is challenging only by measuring structural metrics.

In this study, we used shredder diversity (taxonomic and functional diversity) and OMB rates to quantify the effects of urbanization on stream integrity in an Andean catchment influenced by a rapid and unmanaged metropolitan expansion. In particular, we examined (i) how shredder invertebrates are affected by an urbanization gradient, (ii) whether OMB rates are similarly affected by the urbanization gradient, and (iii) which environmental variables altered by urbanization are disturbing the biodiversity – ecosystem function relationship (i.e. shredder-mediated OMB rates). We hypothesized that the loss of shredder diversity because of urbanization is also propagated in the ecosystem function in Andean streams.

## 2 Material and Methods

### 2.1 Study area and sampling sites

This study was conducted in the city of Loja and its surrounding areas in the southern Ecuadorian Andes (**Figure 1**). The study area, a headwater catchment of the Zamora River basin, is about 276 km^2^ with a population of ≈ 200,000 inhabitants (Iñiguez-Armijos et al., 2014; Chuquimarca et al., 2019). The urban area of the city is highly developed, presenting a mixture of residential and commercial land. The surroundings of the city are mainly covered by pastures land, after pastures, the native forests extend to the catchment divide. Also, a few small plots for farming and small pine plantations can be found in the surroundings of the city. Historically, the central city area has been located along the valley and has grown towards the outskirts. This urban sprawl has transformed land cover around the city from forest to pasture and then to urban over decades leaving an urbanization gradient within the catchment. Sewage is directly discharged to streams and agricultural and urban runoff is not managed Therefore, we selected 12 independent streams along such urbanization gradient, having three replicated streams at urban, pasture, forest mixed with pasture, and forest sites, respectively. At each stream, we placed a sampling site consisted of a 30 m reach located downstream of the target land cover.

**Figure 1.**
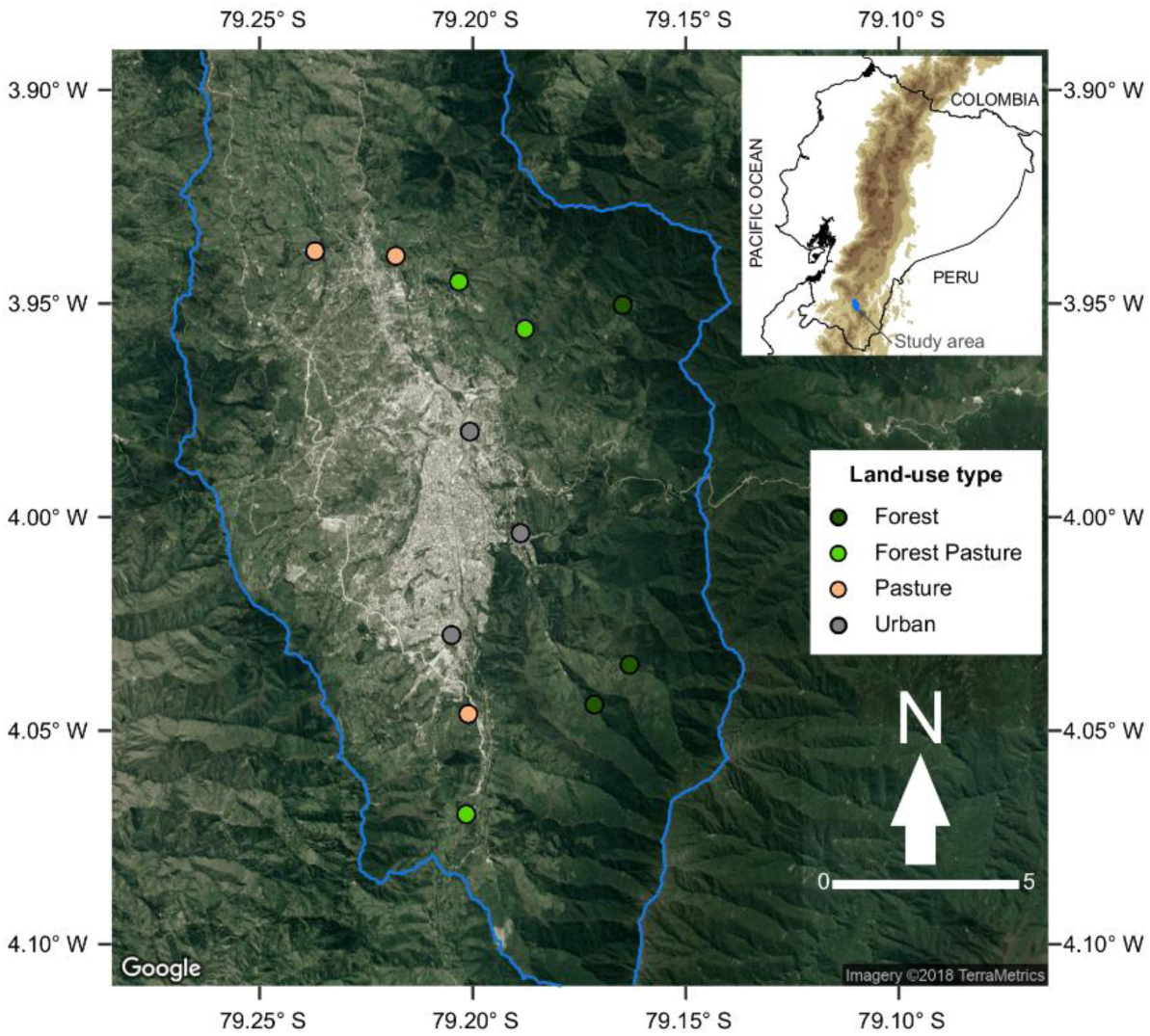
Location of the study area and sampling sites around the city of Loja in the southern Ecuadorian Andes. Colors indicate the land-use gradient studied from forest to urban sites.

### 2.2 Environmental variables

In the dry season of 2016 (September), during the period of stable flow conditions, a total of 16 environmental variables were measured (Table 1). Water temperature, conductivity, pH, and dissolved oxygen were measured using a YSI 556 multi-probe meter (Yellow Springs, Ohio, USA); whereas the concentration of dissolved inorganic nitrogen (DIN = nitrate + nitrite + ammonium), dissolved reactive phosphorus (DRP), and turbidity were determined in laboratory using standard methods (APHA, 2017). Channel width and depth, and current velocity were determined at four to six points along each sampling site using a flowmeter FP311 (Global Water, California, USA). These variables were measured four times in seven-day intervals over throughout the OMB experiment (see below). Substrate size was estimated visually by randomly place 4 plots (1 × 1 m) at each sampling site. The percent of silt, sand, gravel, pebble, cobble, boulder, bedrock was determined according to the Wentworth scale to calculate a substrate index (Harding et al., 2009).

We quantified the percentage of land covered by native forest, pasture, crops, plantations, and urbanization within each catchment of the 12 selected streams. Land cover was obtained upstream the sampling sites using an open-access GIS and a land use map for 2016 obtained in another study (Iñiguez-Armijos pers. comm.). In addition, we quantified riparian canopy cover (%) at four equidistant points from hemispherical photographs taken at 1.3 m above the water surface. Photographs were acquired with a digital camera equipped with a fish-eye lens and then processed in Gap Light Analyzer Version 2.0.

### 2.3 Ecosystem functioning

Leaf-litter breakdown was used as a measure of stream ecosystem functioning. We applied the leaf bag technique (Bärlocher, 2005; Benfield et al., 2017) using freshly fallen leaves of Andean alder (*Alnus acuminata* Kunth), croton (*Croton rimbachii* Croizat), and eucalyptus (*Eucalyptus globulus* Labill.). These species are frequent in the riparian zone of the studied streams, but only croton is a native species whereas alder and eucalyptus have been planted. By using different litter types, we considered potential variation in shredder colonization and breakdown rates. We enclosed 4 ± 0.05 g of air-dried leaves in 15 × 15 cm coarse mesh bags (10 mm) to allow access by shredders. Moreover, fine mesh bash (0.5 mm) were also used to estimate only microbial breakdown by preventing shredders access. This step allowed us to quantify the contribution of shredders to total breakdown (McKie et al., 2009), as explained below.

At each sampling site, four coarse and fine mesh bags of each litter type were tied to the substrate with iron bars and incubated for 30 days before retrieval. Timing was selected to achieve 50% of mass loss based on a previous study (Iñiguez-Armijos et al., 2016), which minimize variation between leaf bags as a result of extended incubation periods (Tiegs et al., 2009). After retrieval, leaf bags were transported individually in zip-lock bags to the laboratory and leaf material was carefully rinsed to remove sediments and recover invertebrates. Then, we dried (48 h/50°C), weighted, combusted (4 h/500°C), and reweighted leaf material to obtain ash-free dry mas (AFDM) (Bärlocher, 2005). An extra set of 5 leaf bags of each litter type was similarly treated to correct mass loss of the final AFDM caused by handling (Bärlocher, 2005).

Over the 30 days, at each sampling site, we recorded water temperature in 1 h intervals using HOBO pendant data loggers (Onset, Massachusetts, USA) placed within the mesh bags and secured to the iron bars. This data was used to calculate the sum of degree days (dd^−1^) during the incubation period to standardize breakdown rate (*k*) by temperature, and to reduce differences between stream types. Then, breakdown rates were estimated using an exponential decay model per degree days (*k* dd^−1^), as follows:

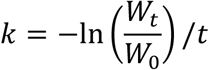

where *W*_0_ is initial air-dried mass, *W*_*t*_ is final AFDM at time *t*, and *t* is incubation time in degree-days. Total breakdown rates (*k*_total_), which is breakdown in coarse mesh bags, were corrected for microbial breakdown (*k*_coarse_ – *k*_fine_) to calculate the contribution of shredders invertebrates (*k*_shredders_) to this ecosystem process (McKie et al., 2009) (Table 1).

### 2.4 Shredder diversity

All invertebrates recovered from the coarse mesh bags were counted and identified to the lowest taxonomic level (usually genus) using available literature for South America (Dominguez and Fernández, 2009). Then, we assigned the invertebrates to a functional feeding group (Ramírez and Gutiérrez-Fonseca, 2014). We only selected shredder invertebrates for this study to be directly associated with leaf litter breakdown. We classified all shredder invertebrates according to three biological traits (size, life cycle duration, respiration) and 14 different modalities based on literature available (Tachet, 2000). Traits such as food, dietary preference, or reproduction were excluded to be general for shredders in this study.

For each litter type, we calculated taxonomic richness (taxa per bag–1) and abundance (individuals per bag–1) of shredders in R environment (R Development Core Team, 2019) using the ‘vegan’ and ‘BiodiversityR’ packages (see Table 1). Also, we estimated functional richness and Rao’s quadratic entropy (RaoQ) based on Gower’s distances of our biological traits using the ‘FD’ package (Laliberté et al., 2014). Particularly, we used these metrics for being adequate to test ecological impairment across environmental gradients. Functional richness measures the amount of multidimensional functional space occupied by the community in multi-trait studies; whereas RaoQ measures the differences among functional entities within a community (Villéger et al., 2008; Laliberté and Legendre, 2010). Due to no shredder species were recorded in the most urbanized streams, for practical and representation purposes we added zero values of functional diversity for these sites.

### 2.5 Urbanization gradient

We produced an urbanization gradient using the environmental variables indicated in Table 1 in a sparse Principal Components Analysis (sPCA) (Zou et al., 2006; Croux et al., 2013), because it facilitates interpretation by reducing the number of variables loading in a principal component. We run the sPCA using the ‘pcaPP’ package (Filzmoser et al., 2018), and following an open access routine for R available at https://github.com/rbslandau/Function_div, which has been used to obtain a similar environmental stress gradient (Voß and Schäfer, 2017). Prior to sPCA, the 16 environmental variables were inspected for normality using the Shapiro-Wil test (‘stats’ package), and were log or square root-transformed if needed. Logit transformation was used for percentages such as canopy cover. To address collinearity, we applied the Variation Inflation Factor (VIF) using the ‘usdm’ package (Naimi, 2015) and considering a VIF threshold of > 7 to exclude variance-inflated variables (Zuur et al., 2007). We used this method because also non-linear relationships can be detected in contrast to correlation analysis (Feld et al., 2016). Consequently, a set of 11 variables was analyzed in the sPCA, obtaining a first principal component (PC1) explaining 53% of total variance between streams and loading 10 out of 11 environmental variables (see **¡Error! No se encuentra el origen de la referencia**.). The scores of the PC1 increased from forested to urbanized streams facilitating environmental interpretation, i.e. increasing scores are related to increasing stress. Therefore, we refer to PC1 as the urbanization gradient hereafter.

### 2.6 Response to urbanization gradient

Zero inflated models (ZIM) were applied to determine how taxonomic and functional diversity of shredder invertebrates are affected by urbanization gradient. Using the ‘glmmTMB’ package (Brooks et al., 2017), ZIMs were build based on Poisson and Negative binomial distributions for taxonomic and functional metric, respectively. Otherwise, generalized additive models (GAMs) with a gaussian distribution were applied to determine if total (*k*_total_) and shredders-mediated (*k*_shredders_) breakdown rates were similarly affected by the urbanization gradient. We used GAMs because they can fit variation in data structures that have a complex non-linear relationship with response variables (Zuur et al., 2007). Model residuals were tested for normality and homoscedasticity assumptions.

Finally, we applied GAMs to identify which environmental variables associated to the urbanization gradient significantly alter the B-EF relationship. Therefore, we did a variable selection from the environmental variables associated to PC1 following a procedure describe in Feld et al. (2016). This step allowed us to select the least statistically redundant predictors and the most ecologically meaningful ones per each group of variables. To represent the B-EF relationship, we used *k*_shredders_ as an indicator of such interaction considering that this rate reveals the role of shredder invertebrates on OMB during the incubation period. Variables were log or square root-transformed prior to analysis if needed. We started with a full model including all predictors until obtaining a simplified model selected with Akaike’s information criterion (AIC) (Zuur et al., 2007; Wood, 2017). GAM models were build using the ‘mgcv’ package (Wood, 2017).

## 3 Results

### 3.1 Shredder diversity and urbanization

A total of 287 shedder invertebrates belonging to nine genera of seven families and four orders were collected across all sites. No shredders were found at urban sites, whereas the lowest shredder richness was observed at pasture sites with seven taxa and the highest at mixed forest-pasture and forest sites with eight taxa each. Moreover, mixed forest-pasture and forest sites had more individual of shredder invertebrates that other sites. ZIM models (**Table 2**), indicated that the effect of the urbanization gradient was negative in both taxonomic richness and abundance of shredders (**Figure** 2A and B). Urbanization also affected negatively the functional richness and RaoQ of shredder invertebrates, but the response of functional diversity was a gradual decrease along the urbanization gradient (**Figure 2**C and D).

**Table 2.**
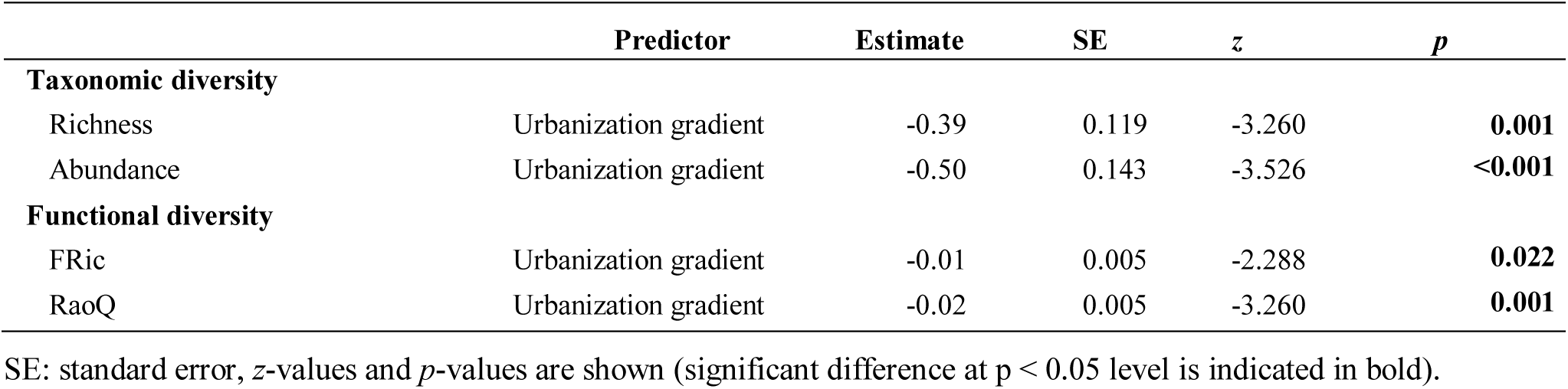
Summary of the zero inflated models (ZIM) showing the response of taxonomic and functional diversity of shredder invertebrates along an urbanization gradient in Andean streams of southern Ecuador.

**Figure 2.**
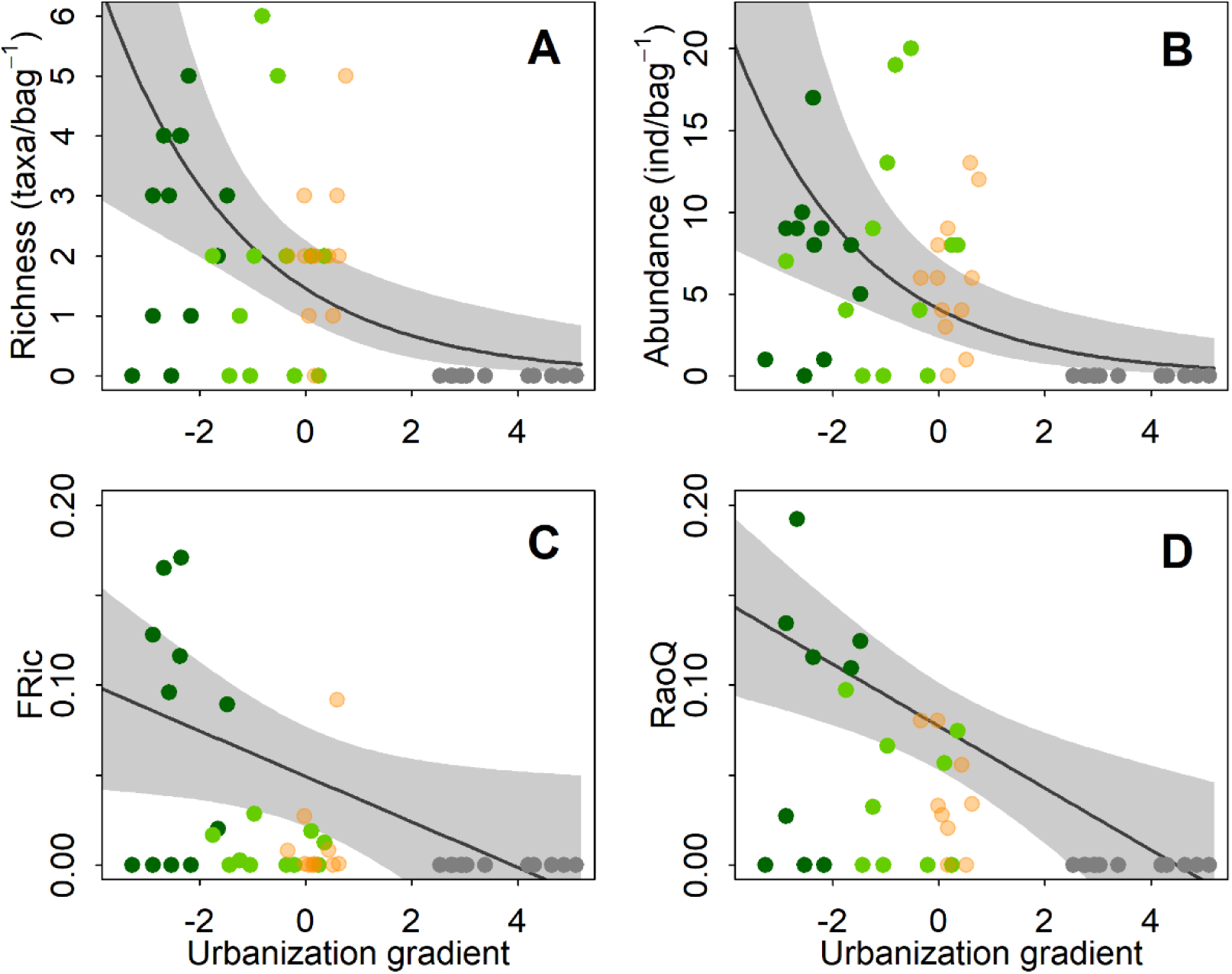
Response of taxonomic richness (A), abundance (B), functional richness (C), and Rao’s quadratic entropy (D) of shredder invertebrates to an urbanization gradient in Andean streams. Gray line represents the ZIM model fitting and the shadows shows 95% confidence interval. Colored circles represent the forest (dark green), forest-pasture (light green), pasture (salmon), and urban (grey) sites, respectively.

### 3.2 Breakdown rates and urbanization

On average, *k*_total_ was 0.0019 dd^−1^ and the contribution of shredders to OMB was much lower than microorganisms (Table 1), indeed *k*_shredders_ represented around 21% of total breakdown. GAM models (**Table 3**), indicated that OMB responded negatively along the urbanization gradient being faster in forest sites diminishing towards urbanized sites (**Figure 3**A). However, the response of OMB mediated by shredders showed a drastic decrease at urban sites (**Figure 3**B).

**Table 3.** Summary of the generalized additive models (GAMs) showing (A) the response of total and shedder-mediated breakdown rates along an urbanization gradient in Andean streams of southern Ecuador; and (B) the response of biodiversity - ecosystem function (B-EF) relationship, represented by *k*_shredders_, to the environmental stressors associated to urbanization that were identified as the best predictors.

|  | Predictor | eDF | F | p | Deviance explained (%) |
| --- | --- | --- | --- | --- | --- |
| **A** |  |  |  |  |  |
| **Breakdown rates** |  |  |  |  |  |
| $k_{\text{total}}$ | Urbanization gradient | 3.491 | 41.02 | **<0.001** | 80.4 |
| $k_{\text{shredders}}$ | Urbanization gradient | 3.151 | 10.19 | **<0.001** | 48.9 |
| **B** |  |  |  |  |  |
| **B-EF relationship** |  |  |  |  |  |
| $k_{\text{shredders}}$ | DRP | 3.005 | 7.28 | **<0.001** | 39.6 |
|  | Substrate size | 1.922 | 12.69 | **<0.001** | 41.4 |

eDF: estimated Degrees of Freedom, $F$ -statistic and $p$ -values are shown (significant difference at $p < 0.05$ level is indicated in bold).

**Figure 3.**
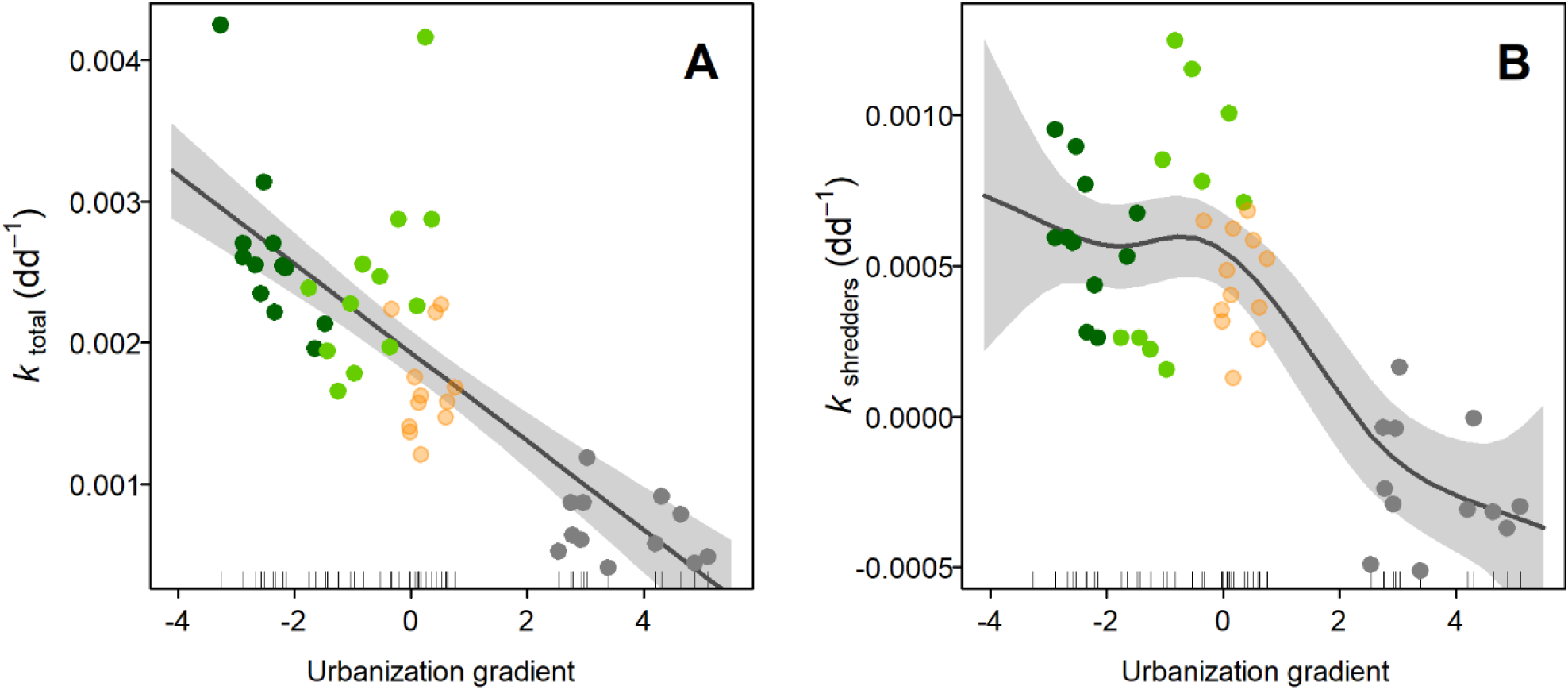
Response of organic matter breakdown to an urbanization gradient in Andean streams. *k*_total_ (A) is breakdown estimated from coarse mesh bags, whereas *k*_shredders_ (B) is the difference of the breakdown estimated between coarse and fine mesh bags. Gray line represents the GAM model fitting and the shadows shows the 95% confidence interval. Colored circles represent the forest (dark green), forest-pasture (light green), pasture (salmon), and urban (grey) sites, respectively.

### 3.3 Urbanization and the biodiversity and ecosystem function relationship

After the selection process of environmental variables associated to the urban gradient (i.e. PC1) for GAM modelling, DRP, pH, turbidity, and substrate size were retained. DIN, and dissolved oxygen were highly correlated with DRP; whereas channel width and depth, and current velocity were highly correlated with substrate size. We excluded urban land cover for being a variance-inflated variable. GAM modeling indicated that DRP and substrate size were the most meaningful variables and had a strong negative effect on *k*_shredders_, i.e. B-EF relationship (**Figure 4**), each explaining 39% and 41% of the deviance explained, respectively (**Table 3**).

**Figure 4.**
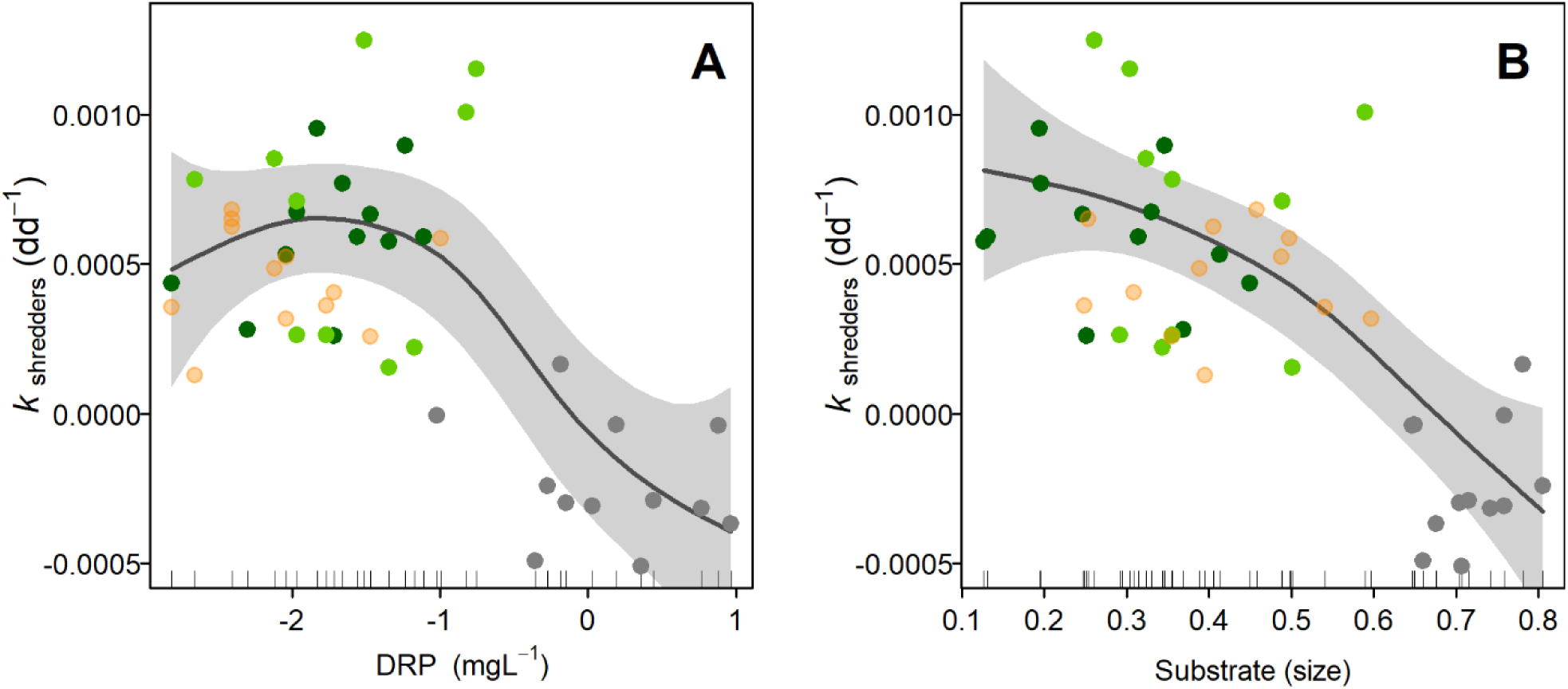
Response of the biodiversity - ecosystem function (B-EF) relationship against two environmental variables associated to the urbanization gradient. The B-EF relationship is represented by *k*_shredders_, while dissolved reactive phosphorus (DRP) and substrate size were selected as the most meaningful variables. Gray line represents the GAM model fitting and the shadows shows the 95% confidence interval. Colored circles represent the forest (dark green), forest-pasture (light green), pasture (salmon), and urban (grey) sites, respectively.

## 4 Discussion

Although the relationship of B-EF has been extensively studied (Tilman et al., 2014), it is still necessary to understand how environmental disturbance gradients can alter this relationship. In this study, we assessed the response of taxonomic and functional diversity of shredder invertebrates as well as OMB rates along an urbanization gradient in Andean streams. We detected that the diversity of shredder invertebrates decreases from natural to more urbanized sites and that shredders are totally absent in urban streams. We also found that OMB rates are negatively affected along the urbanization gradient slowing down this ecosystem process towards urban sites and that urban-associated variables representing water chemistry (e.g. DRP) and physical habitat (e.g. substrate size) of the streams are altering the relationship between shredders and OMB.

DRP is produced by phosphate inputs to stream ecosystems from wastewater coming from sewage systems of the urban areas (Paul and Meyer, 2001). In our study, DRP concentrations were low at forest sites but increased considerably towards urban sites (up to 25-fold greater). At the same time, we observed a decrease in the taxonomic and functional diversity of shredder invertebrates and a slowdown in OMB along the urbanization gradient. Although a moderate increase in the concentration of nutrients such as P can stimulate OMB by shredders in temperate streams (Woodward et al., 2012), in montane Andean streams the increasing of P has been shown to slow down this OMB (Iñiguez-Armijos et al., 2016). Other stream parameters such as dissolved oxygen and DIN were negative and positively associated with DRP. This correlation can be explained by the same land-use intensification along the urbanization gradient, which facilitates the incorporation of organic and inorganic pollutants that interact synergistically and reduce the stream ecosystem integrity (Paul and Meyer, 2001). The increase in N and P concentrations from sewage (e.g. human waste and detergents) decreases the dissolved oxygen (Allan, 2004). Nutrient enrichment favors microbial activity which increases respiration rates and decreases the availability of oxygen in the streams (Walsh et al., 2005). High nutrient concentrations and reduced dissolved oxygen combined can interact reducing or eliminating shredder invertebrates (Couceiro et al., 2006; Iñiguez-Armijos et al., 2016). This pattern is supported by our findings, but we also observed that the reduction in OMB by shredders can be attributed to an abrupt loss of their taxonomic and functional diversity towards more urbanized sites as an effect of nutrient enrichment and low availability of dissolved oxygen.

With regard to substrate size, riparian land-use change and bank erosion produce streambed sedimentation in urbanized streams, reducing substrate size and its heterogeneity affecting negatively the macroinvertebrate diversity (Couceiro et al., 2010). After the loss of macroinvertebrate taxa such as shredders due to the alteration of substrate size and type, their ecological role will be also affected. In our study, we found that the more the urbanization gradient increases, the less the substrate size which is largely dominated by sand, silt and mud. This consequence can explain the strong negative effect of substrate size on OMB mediated by shredders and their loss of taxonomic and functional diversity. Size and type of the substrate condition the presence of several members of the macroinvertebrate community in Andean stream ecosystems as substrate provides habitat, refuge and food (Miserendino and Pizzolon, 2004). Therefore, as the heterogeneity and size of the substrate decrease, there would be a loss taxonomic and functional diversity of shredders invertebrates, and thus their contribution to OMB at the ecosystem level. Other hydraulic parameters such as channel width and depth, and current velocity were negatively correlated to substrate size. In the Andes, streams in natural areas show high current velocity and turbulent flow, they are wide and deep, the streambed presents different substrate sizes and contains several shredder invertebrates taxa (Iñiguez-Armijos et al., 2018a; Vimos-Lojano et al., 2020). On the contrary, the studied streams at urban sites presented a laminar flow as a consequence of sediment deposition in the streambed dominated by small particle sizes such as silt and mud. Also, these streams were less wide due to enclosure, and have been documented as lacking shredder invertebrates (Iñiguez-Armijos et al., 2016). The alteration of these hydraulic variables, as we have described here, has negatively affected the OMB in tropical streams (e.g. Martins et al., 2015); and in our case, we believe that it has resulted in the loss of the taxonomic and functional diversity of shredder invertebrates as well as their capacity for organic matter processing in Andean stream ecosystems.

Decades of research were needed to understand the leading role of biodiversity in the dynamics and functioning of ecosystems, and to quantify the positive effect of species diversity on ecosystem processes (Tilman et al., 2014). Macroinvertebrates play important roles in the functioning of stream ecosystems, among others, shredder invertebrates contribute to the C cycle by feeding on CPOM (Ramírez and Gutiérrez-Fonseca, 2014). However, anthropogenic environmental stress has negatively affected the taxonomic and functional diversity of macroinvertebrates reflecting on the ecosystem functioning of temperate streams (Voß and Schäfer, 2017). In our study on Andean stream ecosystems, we observed that the urbanization gradient notably reduced both taxonomic and functional diversity of shredders which also propagated to OMB rates mediated by this FFG. Indeed, at the most urbanized sites, we did not find shredders indicating a null contribution of macroinvertebrates on organic matter processing in urban streams in the Andes.

The importance of shredders in stream ecosystems is also given as they are key organisms for energy transfer at the low levels of food webs (Ramírez and Gutiérrez-Fonseca, 2014; Merritt et al., 2017). In montane streams the relevance of shredders can be even greater due to leaf litter is the major energy source (Wipfli et al., 2007). When shredders reduce the size of CPOM, they allow other organisms to have nutrients downstream as well as the accomplishment of other ecosystems processes (Woodward, 2009; Vaughn, 2010; Benfield et al., 2017). Our findings suggest that there could be an alteration in the transfer of organic C in urban Andean streams. Nevertheless, more evidence is needed to understand the effect of shedders’ loss on organic C fluxes in these tropical mountain stream ecosystems.

We demonstrate that urbanization affected the B-EF relationships in Andean streams via nutrient enrichment and substrate homogenization causing taxonomic and functional diversity loss of shredders and OMB decline. The loss of keystone species mostly results in a reduction of ecosystem functions (Cardinale et al., 2006; Vaughn, 2010). Also, species loss is associated with the loss of functional biological traits related to ecosystem functioning (Flynn et al., 2009). However, it remains to be understood how biological traits of shredders important for OMB are affected by stressors associated with urbanization in Andean streams. Finally, by losing ecosystem functions, several ecosystem services are also compromised. It is crucial to implement better land-use practices in Andean catchments to guarantee environmental services at lower altitudes.

